# PerturBase: a comprehensive database for single-cell perturbation data analysis and visualization

**DOI:** 10.1101/2024.02.03.578767

**Authors:** Zhiting Wei, Duanmiao Si, Bin Duan, Yicheng Gao, Qian Yu, Ling Guo, Qi Liu

**Affiliations:** State Key Laboratory of Cardiology and Medical Innovation Center, Shanghai East Hospital, Frontier Science Center for Stem Cell Research, Bioinformatics Department, School of Life Sciences and Technology, Tongji University, Shanghai 200092, China; Key Laboratory of Spine and Spinal Cord Injury Repair and Regeneration (Tongji University), Ministry of Education, Orthopaedic Department of Tongji Hospital, Frontier Science Center for Stem Cell Research, Bioinformatics Department, School of Life Sciences and Technology, Tongji University, Shanghai 200092, China; Zhejiang Lab, Hangzhou 311121, China; Shanghai Research Institute for Intelligent Autonomous Systems 201804, China

**Author notes:** Corresponding authors: Correspondence to Qi Liu^1, 2, 3, 4, #^ with, Correspondence to Ling Guo^3, #^ with. These authors contribute equally to this work (Zhiting Wei, Duanmiao Si, Bin Duan, Yicheng Gao).

## Abstract

Single-cell perturbation sequencing techniques (scPerturbation), represented by single cell genetic perturbation sequencing (e.g., Perturb-seq) and single cell chemical perturbation sequencing (e.g., sci-Plex), result from the integration of single-cell toolkits with conventional bulk screening methods. These innovative sequencing techniques empower researchers to dissect perturbation functions and mechanisms in complex biological systems at an unprecedented resolution. Despite these advancements, a notable gap exists in the availability of a dedicated database for exploring and querying scPerturbation data. To address this gap and facilitate seamless data sharing for researchers, we present PerturBase—the first and most comprehensive database designed for the analysis and visualization of scPerturbation data (http://www.perturbase.cn/). PerturBase consolidates 122 datasets from 46 publicly accessible research studies, covering 115 single-modal and 7 multi-modal datasets that include 24254 genetic and 230 chemical perturbations from about 6 million cells. The database provides insights through various software-analyzed results, encompassing quality control, denoising, differential expression gene analysis, perturbation function analysis, and correlation characterization between perturbations. All datasets and in-depth analyses are presented in user-friendly, easy-to-browse pages and can be visualized through intuitive tables and various image formats. In summary, PerturBase stands as a pioneering high-content database, intended for searching, visualizing, and analyzing scPerturbation datasets, contributing to an enhanced understanding of perturbation functions and mechanisms.

**Graphical abstract:** 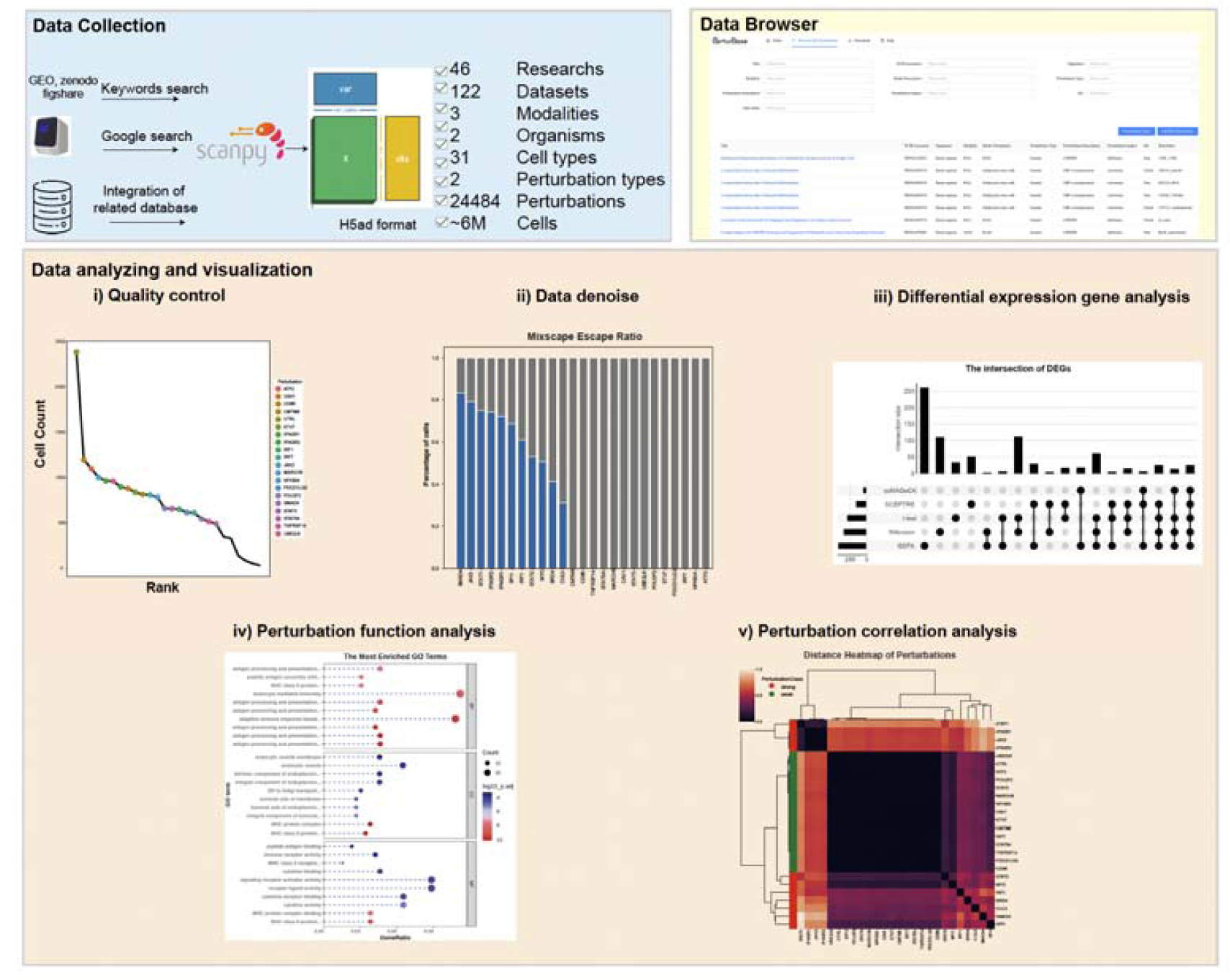

## Introduction

Perturbation experiments explore cellular responses to environmental changes, which can be broadly categorized into two classes: genetic perturbation and chemical perturbation. The perturbation-based omics has become a powerful tool for studying gene functions and a cornerstone of the pharmaceutical drug discovery pipeline(1–3). For instance, library of integrated network-based cellular signatures (LINCS) piloted by Broad Institute has become the rich data sources for genetic and chemical perturbation, and clustered regularly interspaced short palindromic repeats (CRISPR) screening techniques have been broadly used for drug target identification and drug resistance research(4–6). Nonetheless, conventional perturbation technology has at least two major limitations. First, the readout of most are restricted to gross cellular phenotypes, e.g., proliferation, morphology or a highly specific molecular readout. Second, even when in conjunction with more comprehensive molecular phenotyping such as next generation sequencing (NGS), a limitation of bulk assays is that cells ostensibly of the same “type” can exhibit heterogeneous responses(7–9).

Single-cell transcriptome sequencing (scRNA-seq) represents a form of high-content molecular phenotyping that, when combined with conventional perturbation technology, can overcome both limitations. In 2016, single cell CRISPR (scCRISPR) that coupled CRISPR screening and scRNA-seq to enable pooled genetic screens at large-scale single-cell resolution was developed(10–12). The key technical innovation of scCRISPR is the creative design of the lentiviral vector to allow the identification of sgRNA in each cell from sequencing. Meanwhile In 2019, sci-Plex, a method that coupled chemical screening with scRNA-seq to cost-effectively quantify transcriptional responses to hundreds of chemicals in parallel, was proposed by Trapnell et al(9). In contrast to traditional perturbation screening, scPerturbation allows obtaining high-content phenotypes, thus facilitate dissection of complex mechanisms of genes and chemicals in heterogeneous cell populations.

Currently, numerous alternative scPerturbation platforms have emerged. Based on the readout omics, these platforms can be classified into four primary categories: transcriptome-based, epigenome-based, proteome-based and imaging-based. The mainstream scPerturbation platforms are transcriptome-based that combine screens with scRNA-seq, such as Perturb-seq and sci-Plex(9–13). Transcriptome-based platforms have evolved rapidly, with innovations like CROP-seq optimizing the Perturb-seq vector design, reducing the complexity and cost(14). Genome-scale Perturb-seq, introduced by Replogle et al., enables unbiased and comprehensive profiling of genome-scale genetic perturbations affecting 9867 genes(15). What’s more, by applying the technique to multi-omics simultaneously, multi-modal scPerturbation was developed. In 2019, Rubin et al. developed an epigenome-based scPerturbation, named Perturb-ATAC that combines CRISPR interference or knockout with chromatin accessibility profiling in single cells based on simultaneous detection of CRISPR guide RNAs and open chromatin sites by assay of transposase-accessible chromatin with sequencing (ATAC-seq) (16). Mimitou et al. developed ECCITE-seq, which allowed simultaneous detection of transcriptomes, proteins, clonotypes, and CRISPR perturbations from every single cell(17). Recently, the new multi-modal scPerturbation platforms called Perturb-map and Perturb-FISH which combines CRISPR with imaging and spatial transcriptomics were developed to enable the identification of genetic determinants of tumor composition, organization and immunity(13,18).

The analysis of scPerturbation data is a major challenge due to its inherent noise. Thus, a series of specialized bioinformatic tools have been developed. Generally, these tools focus on three parts: (1) Data preprocessing, including basic scRNA-seq quality control and normalization, such as MIMOSCA, MUSIC, and SCREE(10,19,20). (2) Data denoising, including single-cell imputation, escaping cells filtering, and latent component factors decomposing, such as MUSIC, mixscape, and GSFA(19,21,22). (3) Functional analysis, including prioritizing the impact of each perturbation, identifying the function of each perturbation, inferring regulatory network and gene interaction, such as MUSIC, Normalisr, scMAGeCK, Pando, GSFA and SCEPTRE(19,22–25).

scPerturbation is widely applied in various fields due to its powerful capabilities, including linking genotype to phenotype(11,12,15), dissecting genetic regulations, and deciphering drug mechanism. For example, Jaitin et al. revealed the effect of 22 TFs on the regulation of antiviral, inflammatory, or developmental processes in Lipopolysaccharide (LPS) stimulated born marrow cells (BMCs) by CRISP-seq(12). Perturb-ATAC, Spear-ATAC, and CRISPR-sciATAC could revealed epigenetic landscape remodelers in human B lymphocytes and leukemia cells(26,27). Using Perturb-map, Dhainaut et al. discovered that the knockout of tgfbr2 in lung cancer cells promotes tumor microenvironment remodeling and immune exclusion(18). Based on sci-Plex, Trapnell reveal substantial intercellular heterogeneity in response to specific chemicals and find that the main transcriptional responses to HDAC inhibitors involve cell-cycle arrest(9).

Despite the widespread use of scPerturbation, a significant gap remains in the availability of a dedicated database for exploring and querying scPerturbation data. We noticed that recently scPerturb is developed for scPerturbation data exploration, however, its utility is constrained by the absence of dedicated features for querying, visualizing and further interpreting the data(28). To this end, we introduce PerturBase, the first and most comprehensive database that integrates 122 scPerturbation datasets from 46 publicly accessible research studies. The molecular readouts of these datasets comprise 115 single-modal and 7 multi-modal datasets. Among these, 101 datasets have been subjected to genetic perturbations, while the remaining 21 were influenced by chemical perturbations. Over 90% of the datasets are derived from Homo sapiens studies. PerturBase features two modules: the dataset browser module, and the dataset analyzing module. The dataset browser module facilitates streamlined exploration of all 122 datasets, offering filters by organism, modality, perturbation type, perturbation name, and perturbation function. The dataset analyzing module provides insights into a range of software-analyzed results, primarily including: (1) quality control outcomes; (2) denoising results; (3) differential expression gene (DEG) analysis; (4) perturbation function analysis; and (5) characterizing the correlation between perturbations. In summary, PerturBase stands as the pioneering high-content database, designed for the searching, visualization, and analysis of scPerturbation data. Its extensive data repository and diverse functionalities make it an indispensable resource in the scPerturbation research community.

## Materials and methods

### Data collection

In our current study, the scPerturbation datasets were primarily obtained through three methods (Figure 1A). (1) We collected the scPerturbation data through a large-scale search in Gene Expression Omnibus, Zenodo (https://zenodo.org), Figshare (https://figshare.com) using keywords such as “perturb seq”, “high content crispr screening”, “single cell crispr screening” and “single cell perturbation”; (2) We selected 10 representative scPerturbation platforms in scPerturbation field and obtained articles citing the aforementioned platforms through Google. The “BioProject” ID of the articles were retrieved via NCBI’s eSearch Application Programming Interface (API). Subsequently, we manually confirmed the presence of scPerturbation data from the identified “BioProject” ID; (3) scPerturbation data mentioned in scPerturb and sc-pert were also maintained in PerturBase(28,29). In summary, the current version of PerturBase contains 122 scPerturbation datasets (Figure 1A, Supplementary table S1). In terms of perturbation type, the collection encompasses 24254 genetic and 230 chemical perturbations (Figure 1B); In terms of perturbation modality, it covers 115 single-modal data and 7 multi-modal data; In terms of species, it contains Homo sapiens and Mus musculus. Notably, most perturbations are predominantly applied in a single dataset, particularly in the case of genetic perturbation (Figure 1B). The total number of cells per dataset is usually restricted by experimental limitations, though has increased over time (Figure 1B, Supplementary Figure S1). Therefore, there is a tradeoff between the number of perturbations and the mean cells per perturbation in a dataset (Supplementary Figure S1).

**Figure 1.**
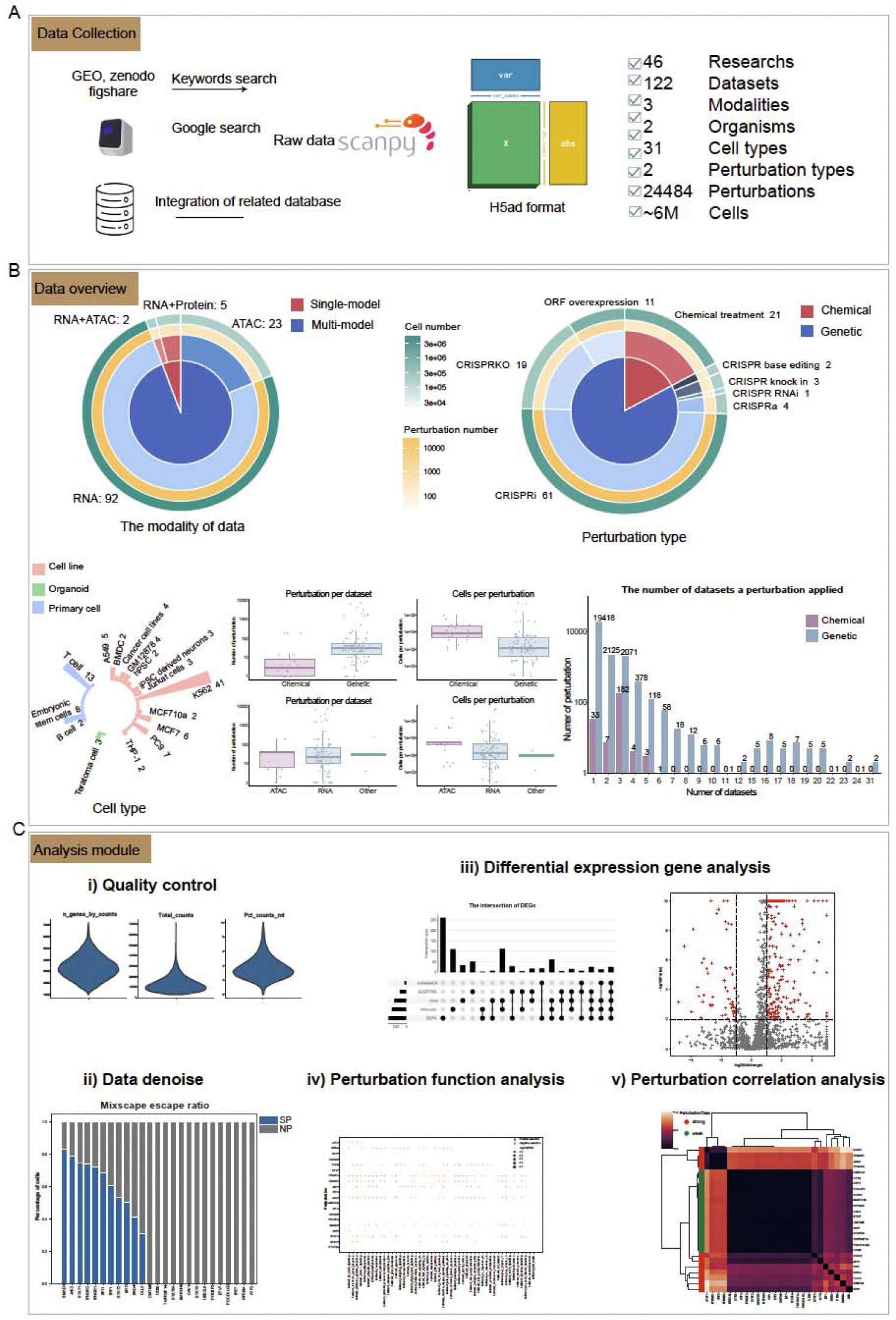
Overview of the design, statistics, and functions of PerturBase. (A) Data collection, construction and summary of the PerturBase resource. (B) Overview of modalities, perturbation types, and cell types in PerturBase. (C) The main functions in PerturBase.

### Processing of the scPerturbation RNA-seq data

#### 1: The assignment of perturbation to a cell

For scCRISPR data, we allocated guide RNAs (gRNAs or sgRNAs, indicating the targeted gene in each cell) to a cell based on two criteria. (1) For cells with sgRNA information already provided in the original research, we used these results directly; (2) For cells where the original research included sgRNAs count matrix data, we allocated sgRNAs using a threshold strategy similar to Mixscape, specifically considering a sgRNA valid if it had a minimum of 5 counts(21). Cells without any sgRNA induction were marked as blank controls, and those with non-targeting sgRNAs (e.g., Green Fluorescent Protein, GFP) were labeled as “CTRL”. For single cell chemical perturbation data, we used label information provided in the original research.

#### 2: Data quality control

For each scPerturbation data, we utilized the anndata and Scanpy Python package to uniformly process it into the h5ad format(30). (1) cells with fewer than 200 expressed gene, classified as blank control, or containing a large fraction of mitochondrial genes (over 10%) were filtered; (2) genes expressed in less than 3 cells were filtered. (3) Datlinger et al. concluded that at least 30 cells are required to capture each perturbation phenotype(14). Therefore, the perturbations except negative control with perturbed cells lower than 30 (default) were not considered in PerturBase. It should be noted that if no perturbation meets these criteria, the dataset is not subjected to further analysis; (4) PerturBase adopts the global scaling normalization method in Scanpy to scale the expression in each cell to 10,000, followed by logarithmic transformation; (5) after normalization, PerturBase adopts “highly variable genes” with default parameter in Scanpy to identify highly variable features for scPerturbation data. To strike a balance between computational efficiency and data retention, PerturBase maintains a gene count ranging from a minimum of 2000 to a maximum of 4000 genes; (6) PerturBase performs principal component analysis (PCA) on the top variable genes, adhering to default parameters (n_components = 50) to reduce dataset dimensionality; (7) After dimension reduction, PerturBase performs clustering based on the leiden algorithm with default parameters; (8) For user convenience, we employed clusterProfiler to obtain gene symbol, Entrez and Ensembl IDs for genes in each dataset, making them readily accessible to users within their dataset of interest(31).

### Processing of the scPerturbation ATAC-seq data

#### 1: Guide RNA assignment

The procedures parallel those of the scPerturbation RNA-seq in guide RNA assignment.

#### 2: Peak count matrix generation

scPerturbation ATAC-seq data exists in two primary formats: the Peak Count Matrix and the 10x Fragment TSV file. In the case of the Peak Count Matrix file, its raw data was preserved. As for the 10x Fragment TSV file, we utilized macs2 within the Signac CallPeaks function to derive the Peak Count Matrix(32,33).

#### 3: Data quality control

For each scPerturbation ATAC-seq dataset, we consistently processed its Peak Count Matrix into the h5ad format using the Seurat and Signac R packages. (1) filtering cells with expressed peak numbers fewer than 200 or more than 30000, fragment ratio in peak less than 0.15, blacklist fraction greater than 0.05, nucleosome signal exceeding 4, transcription start site (TSS) enrichment below 4, or being a blank control; (2) filtering peaks expressed in fewer than 10 cells; (3) The perturbations except negative control with perturbed cells lower than 30 (default) were not considered; (4) scaling the peak count matrix using the frequency-inverse document frequency (TF-IDF) method in Signac. Subsequent procedures align with those employed for scPerturbation RNA-seq in data quality control from step (5).

### Data denoising of the scPerturbation RNA-seq

Alternative sources of variation, including transduction replicate identity, cell cycle stage, and activation of cellular stress responses, can confound the downstream analysis of scPerturbation data. To mitigate these issues, we employ Mixscape(21) from pertpy to compute the local perturbation signature for each cell (Figure 1C). The core concept is to isolate the genetic perturbation’s effect by subtracting the averaged expression of the K nearest cells from the negative control pool from each cell. Consequently, Mixscape extracts the component of the cell’s profile that solely reflects the genetic perturbation. As recommended, we set the number of neighbors K to 20. Following the denoising of the scPerturbation log-transformed data, we conduct dimension reduction and clustering analysis in line with our established protocols.

### The identification of escaping cell after denoising

The evaluation of sgRNA knockout efficiency in genetic screening and off-target effects in chemical screening is crucial(9,34). In genetic screening, sgRNAs direct Cas9 to specific genomic loci, yet only about 70–80% effectively induce the desired impact on the targeted gene. This indicates that in 20–30% of cells harboring an identified sgRNA, the gene may remain unaffected or partially impacted, exhibiting a wild-type phenotype (defined as “escaping” cell). Similar to genetic screening, chemical screening also exhibit off-target effects(9). Such occurrences can skew the assessment of a perturbation’s effect. Consequently, a filtering step to eliminate these cells is necessary. Mixscape leverages the cell’s transcriptome as a phenotypic indicator of the perturbation’s impact and has devised a method to systematically identify and exclude escaping cells. Mixscape’s fundamental premise is that each perturbation class comprises a mix of two Gaussian distributions: one representing successful perturbed (SP) and the other non-perturbed (NP) cells. The transcriptional profile distribution of NP cells should align with control (CTRL) cells. Mixscape computes the posterior probability of a cell belonging to the SP class, and categorize those with a probability over 0.5 as SP cells. This approach, applied across all perturbations, enables the identification of all SP cells and assesses the targeting efficacy of genetic and chemical perturbation. Notably In our study, we conducted additional analysis. We postulated that the targeting efficiency of a perturbation would not be less than 20%. Hence, if the SP / (NP + SP) ratio for a perturbation is below 20%, it suggests that the perturbation is truly unlikely to induce a significant phenotypic change in cells. Such perturbations are deemed “weak” perturbations and all corresponding cells are retained for analysis(21). Conversely, if the SP / (NP + SP) ratio exceeds 20%, the perturbations are categorized as “strong” perturbations, and NP cells are filtered out, adhering to Mixscape’s criteria. Following the denoising and exclusion of NP cells, we conducted dimension reduction and clustering analysis on the log-transformed scPerturbation data, as previously described.

### The identification of differentially expressed genes of a perturbation

In our current version of PerturBase, we employed five methods to detect differentially expressed genes (DEGs, Figure 1C), including three specifically developed for scPerturbation (scMAGeCK, GSFA, and SCEPTRE) and two being commonly used methods in scRNA-seq provided by Scanpy (Wilcoxon and t-test). For scPerturbation ATAC-seq data, three methods provided by Seurat were employed (Wilcoxon, t-test and logistic regression).

For scMAGeCK, genes with regulatory score greater than 0.2 or less than −0.2 and *P*-value lower than 0.05 are defined as DEGs. For GSFA, genes with *P*-value lower than 0.05 are defined as DEGs. For SCEPTRE, genes with foldchange greater than 2 or less than 0.5 and *P*-value lower than 0.05 are defined as DEGs. For Wilcoxon and t-test, genes with foldchange greater than 2 or less than 0.5 and *P*-value lower than 0.05 are defined as DEGs. The input for the five methods is the expression profile of highly variable genes. All the parameters in the above five methods were set as default. For logistic regression (LR), genes with foldchange greater than 2 or less than 0.5 and *P*-value lower than 0.05 are defined as DEGs.

### The evaluation of the function of a perturbation

We evaluated the function of a perturbation through three distinct methodologies (Figure 1C): (1) by the enrichment analysis of a perturbation’s DEGs. PerturBase performs gene ontology (GO) and Kyoto Encyclopedia of Genes and Genomes (KEGG) enrichment analysis for each perturbation using the DEGs of each perturbation. PerturBase provides detailed enrichment results for each perturbation, including both individual and aggregated analyses. Specifically, we present individual enrichment outcomes for each perturbation and a collective analysis for the top 25 most impactful perturbations. Enrichment terms are defined as significant if they exhibit a Q-value below 0.05 and a *P*-value under 0.01; (2) by characterizing a perturbation’s association with the MSigDB signatures (gene sets). We utilized the RRA module of scMAGeCK to link perturbation with MSigDB signatures(24). At the current version of PerturBase, 50 well defined hallmark signatures in MSigDB were downloaded for analysis. A perturbation is considered to significantly negatively regulate a phenotype corresponding to a signature if the “FDR.low” value is under 0.01. Conversely, it is deemed to significantly positively regulate a phenotype if the “FDR.high” value is below 0.01; (3) by the evaluation of a perturbation’s clustering distribution(35). This analysis consists of two parts: firstly, we assess whether a perturbation is preferentially enriched in a specific cluster compared to the CTRL. Secondly, we evaluate whether its distribution across clusters significantly deviates from that of the CTRL. This evaluation utilizes the chi-squared test, considering a perturbation to significantly impact cell behavior if the *P*-value is 0.01 or lower.

### Characterizing the correlation between perturbations

Perturbations with shared mechanisms or target tend to produce similar shifts in expression profiles. Therefore, by characterizing the correlation between perturbations using expression profiles, we can describe the difference or similarity between perturbations in terms of mechanism or perturbation target. In our current study, we characterize the correlation between perturbation with three methods (Figure 1C): (1) using processed expression profile. Firstly, the mean expression profiles of the perturbations in a dataset were calculated. Then, the correlations between perturbations were calculated using cosine similarity; (2) using E-distance function in pertpy(28). E-distance is a statistical distance measure which compares the mean pairwise distance of cells across two different perturbations to the mean pairwise distance of cells within the two distributions. A large E-distance of perturbed cells from unperturbed indicates a strong change in molecular profile induced by the perturbation. Similar to Replogle et al., we compute the E-distance after PCA; (3) characterizing the correlation between perturbations using latent factors output by GSFA(22). GSFA describes the effects of a perturbation through a set of latent factors which represent biological pathways or functional units. The correlations between perturbations were calculated using cosine similarity with the latent factors of perturbations.

### Database construction

PerturBase was built on a Linux server. The web services have been built using Nginx (version 1.24.0). The front end of PerturBase is built with HTML5, JavaScript, CSS and React (version 18.0.0) that consists of the react UI library ant-design (version 4.20.7). All data in PerturBase were stored and managed by MySQL (version 8.0.36). PerturBase has been tested on a number of popular web browsers, including the Google Chrome, Firefox and Apple Safari web browsers. No registration or login is required.

## Results

### Overview of PerturBase

PerturBase curates 122 scPerturbation datasets from 46 publicly research studies, consisting of 115 single-modal data and 7 multi-modal data, covering Homo sapiens and Mus musculus. Of these datasets, 101 datasets were perturbed using 24254 genetic and 21 datasets perturbed with 230 chemical compounds. PerturBase features two distinct modules: the dataset browser module, and the dataset analyzing module. The dataset browser module facilitates streamlined exploration of all 122 datasets, offering filters by organism, modality, perturbation type, perturbation name, and perturbation function (Figure 2A). The dataset analyzing module provides insights into a range of software-analyzed results, primarily including: (1) quality control; (2) denoising; (3) DEGs analysis; (4) perturbation function analysis; and (5) characterizing the correlation between perturbations (Figure 2B-F).

**Figure 2.**
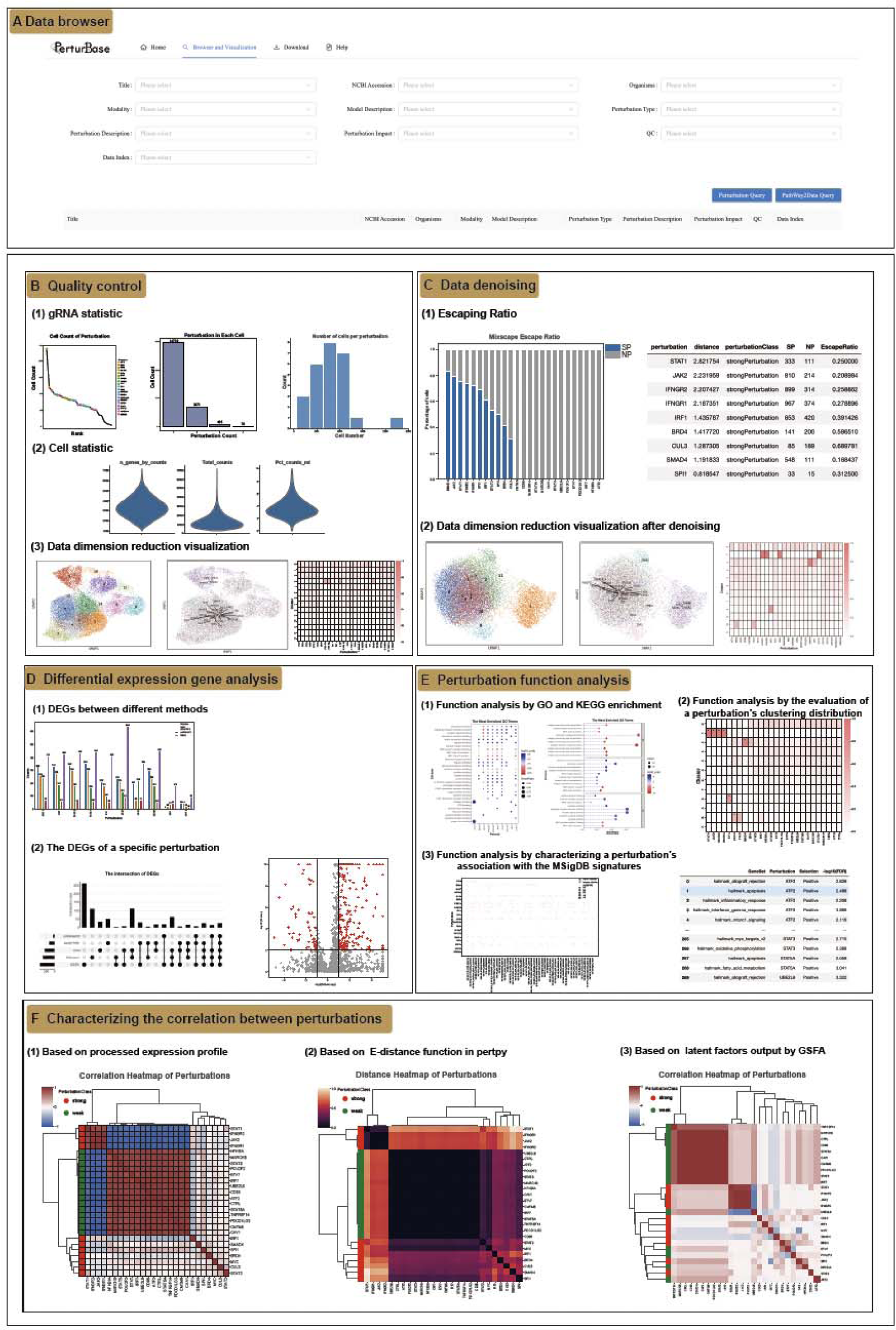
User interface of PerturBase. (A) Data browser module allow users to obtain scPerturbation data base on research title, perturbation, perturbation type, cell line, and etc. (B) Quality control section demonstrates the basic information about a preprocessed dataset. (C) PerturBase adopts Mixscape to alleviate confound factors and Data denoise section shows the escaping ratio and dimension reduction results after data denoising. (D) PerturBase implemented five distinct methods for detecting DEGs in scPerturbation RNA-seq data and three methods for ATAC-seq data. This section offers the DEGs of a perturbation under a chosen method. (E) Function analysis section employed three methodologies to evaluate the functional consequences of perturbations. (F) Correlation section offers the correlations between perturbations.

### The dataset browser module

The dataset browser module provides easy exploration and accession of all 122 perturbed datasets with 12 keywords (Figure 2A). For instance, the “Perturbation” keyword describes the perturbations a dataset maintained and users can conveniently search datasets contains the perturbation of interest. The “PathWay2Data” keyword describes the functions of a perturbation and uses can search datasets contain the perturbations that have the function of interest (Materials and methods). Upon selecting a dataset of interest, users can access a wealth of detailed information and an array of downstream analysis results related to that dataset.

### The dataset analyzing module

The dataset analyzing module of PerturBase primarily encompassing five components: (1) the visualization of quality control result; (2) the visualization of data denoise result; (3) the visualization of perturbations’ DEGs; (4) the visualization of perturbations’ functions; (5) the visualization of perturbations’ correlation (Figure 2B-F).

### The visualization of quality control result

The quality control results demonstrate the basic information about a preprocessed dataset such as the perturbations the dataset contains and the distribution of cell numbers of each perturbation (Figure 2B). The “Perturbation in Each cell” describes the number of perturbations a cell accept. This information is useful if users want to access data that contain combination perturbations. We utilized UMAP for visualizing the clustering results, and employed heatmaps to illustrate the distribution of each perturbation across the clusters. In summary, the quality control results provide detailed information about the dataset after quality control.

### The visualization of data denoise result

Alternative sources of variation, including transduction replicate identity, cell cycle stage and the activation of cellular stress responses would confound downstream analysis. Therefore, PerturBase adopt Mixscape(21) of pertpy to alleviate those confound factors by calculating the local perturbation signature for each cell (Figure 2C, Materials and methods). In addition, we utilized UMAP and heatmap to visualize and compare the clustering results before and after denoising. After denoising, we further employed Mixscape to identify escaping cells. The escaping cell was defined as a cell although receives perturbation but do not exhibit the expected phenotype due to off-target effects. Those escaping cells will influence the estimation of the effect of a perturbation. Therefore, Mixscape was employed to identified those escaping cells while evaluating the efficacy of each perturbation (Materials and methods). In our current study, a further analysis was performed. We utilized the efficacy information to classified a perturbation to “weak” perturbation or “strong” perturbation(21). The strong perturbation usually has a great impact on the cell’s phenotype while the weak perturbation has a little impact on the cell’s phenotype. This information provides an insight into the function of a particular perturbation, which may aid our understanding of the roles of a perturbation. All the above information including the results output by Mixscape and filtered dataset can be browsed and accessed in our current version of PerturBase.

### The visualization of perturbations’ DEGs

In our study, we implemented five distinct methods to detect DEGs for scPerturbation RNA-seq data(22,23,30) and three methods for scPerturbation ATAC-seq data(32,33), as detailed in the Materials and Methods section (Figure 2D). To showcase the number of DEGs identified by each of the five software tools, we used barplot to effectively highlighting the variability in perturbation detection. By default, the analysis focuses on the top 25 perturbations with the most pronounced effects. However, if users have interest in a particular perturbation, they can delve deeper into the associated DEGs. This is facilitated through interactive tables and volcano plots, which become available once a specific perturbation and software tool are selected. Additionally, PerturBase incorporates the UpSet R package to illustrate the overlaps and distinctions in DEG identification across different software tools for a given perturbation. This visualization aids in understanding the consensus and discrepancies in DEG identification among various analytical approaches.

### The visualization of perturbations’ functions

As detailed in the Materials and Methods section, to thoroughly assess the functional consequences of a perturbation, we employed three distinct methodologies (Figure 2E). The outcomes of these analyses are effectively visualized using barplots and heatmaps. By design, our interface prioritizes and displays only the top 25 perturbations, identified as having the most significant effects. In visualization of the GO and KEGG enrichment results, we meticulously present both individual and aggregated enrichment analyses for each perturbation, with a focused review of the top 25 perturbations that exhibit the most pronounced impact. Additionally, PerturBase enhances user engagement by allowing the exploration of enrichment results for differentially expressed genes (DEGs) identified through specific software tools, tailored to individual perturbations.

### The visualization of perturbations’ correlation

Perturbations sharing similar mechanisms or targets often manifest comparable shifts in gene expression profiles. Consequently, by examining these expression profiles, we can elucidate the differences or parallels between perturbations in terms of their underlying mechanisms or specific targets. In our study, we approach the characterization of these correlations through three distinct methods (Figure 2F): (1) analyzing the correlation between perturbations based on processed expression profiles; (2) employing the E-distance function in pertpy to quantify the relationship between perturbations; (3) leveraging latent factors derived from GSFA to further interpret these correlations. The findings from these analyses are concisely presented in a heatmap format, facilitating an intuitive understanding of the relationships between various perturbations.

## Case study

The programmed death-ligand (PD-L)1 is frequently observed in human cancers and can lead to the suppression of T cell-mediated immune responses. To demonstrate the capabilities of PerturBase, we utilized a scPerturbation dataset from Papalexi et al. to investigate the regulatory mechanisms of the expression of PD-L1(21). This comprehensive analysis highlights the efficacy of PerturBase in gene function research. The dataset encompasses 25 perturbations, with cell counts ranging from 33 to 1197 post-quality control, each subjected to a unique genetic perturbation (Figure 3A). Of these, 11 were categorized as “strong” perturbations, while the remaining 14 were deemed “weak” perturbations. The sgRNA efficiencies in the strong perturbations varied from approximately 50% to 80%, suggesting that some cells, despite receiving an sgRNA, did not exhibit the expected phenotype, thus being classified as “escaping cells” (refer to Materials and methods for details). For instance, as depicted in Figure 3B, the identified escaping cells of IFNGR1 show a wild-phenotype and IFNGR1’s target gene PD-L1 was not affected. After filtering out escaping cells, two clear groups of cells were observed, including a cluster consisting of IFNGR1, IFNGR2, JAK2, STAT1 and a second cluster consisting of IRF1, underscoring the necessity and effectiveness of the filtering strategy employed by PerturBase.

**Figure 3.**
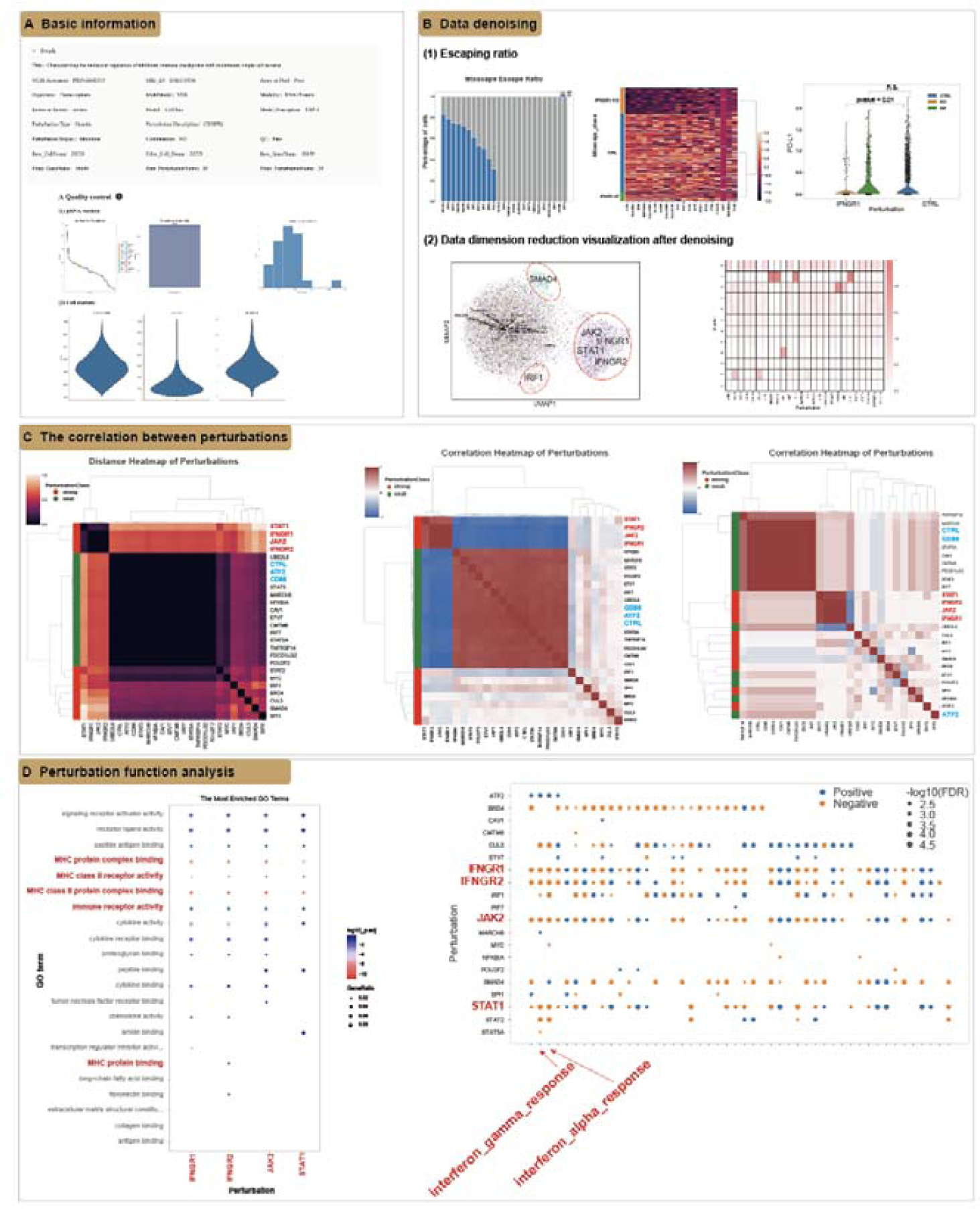
Case study of a scPerturbation dataset to demonstrate the capabilities of PerturBase. (A) The basic information of the selected dataset after quality control. (B) The escaping cells show a wild-type phenotype and after filtering out escaping cells, two clear groups of cells were observed, underscoring the necessity and effectiveness of the filtering strategy employed by PerturBase. (C) IFNGR1, IFNGR2, JAK2, and STAT1 perturbations have strong correlations, suggesting their shared functions, which is concordant with prior knowledge. (D) IFNGR1, IFNGR2, JAK2, and STAT1 perturbations leads to the inhibition of immune response pathway, culminating in the reduced expression of PD-L1.

As Figure 3C shows, IFNGR1, IFNGR2, JAK2, and STAT1 have strong correlations, which is concordant with prior knowledge(36,37). Additionally, perturbations such as STAT3, CD86, and ATF2 exhibit a high degree of similarity to the CTRL group, suggesting minimal phenotypic impact on the cells. This observation is consistent with their previous classification as “weak” perturbations. The DEGs analysis results shows that IFNGR1, IFNGR2, JAK2, STAT1, and IRF1 are the positive regulators of PD-L1(21). Further investigation into the mechanisms driving PD-L1 down-regulation revealed that these four perturbations are central to the immune response pathway, particularly in the context of interferon (IFN)-γ signaling. The disruption of these components consequently leads to the inhibition of this pathway, culminating in the reduced expression of PD-L1. In conclusion, these insights underscore the comprehensive nature of PerturBase as a scPerturbation database. It stands out as an indispensable tool for researchers seeking to access and analyze perturbation data, providing a robust platform for the exploration of perturbation functions and mechanisms.

## Conclusions and future development

PerturBase provides two major modules covering dataset searching and analyzing for understanding of the high-content scPerturbation datasets. The dataset browser module provides easy exploration and accession of all 122 perturbed datasets with 12 keywords. To enhance the interpretability of high-content perturbation resources, the dataset analyzing module provides visualization for five major analytic results that cover the primary requirements, including the quality control, denoise, identification of DEGs, perturbation function analysis, and the correlation between perturbations. Compared to scPerturb, PerturBase offers modules for querying, analyzing and visualizing a more comprehensive scPerturbation data through an interactive interface(28).

Moving forward, we aim to enhance PerturBase in several key areas: Firstly, our ongoing commitment involves the continual curation of datasets, expanding our repository with the latest high-content scPerturbation research, particularly focusing on chemical screening and multi-modal experiments. Secondly, despite providing five analytical results in our dataset analyzing module, we recognize the need for more interactive visualizations for each result. To address this, we are dedicated to developing a more user-friendly and interactive platform in our forthcoming version, facilitating easier access to comprehensive information. Lastly, we plan to incorporate additional analytical modules to deepen the understanding of scPerturbation datasets. In essence, PerturBase stands as the pioneering high-content screening database, specifically designed for the efficient search, visualization, and analysis of scPerturbation datasets. We envision PerturBase becoming an indispensable resource in the field, offering an extensive range of data and functionalities.

## Data availability

PerturBase is an open resource for interactively visualizing and analyzing the comprehensive scPerturbation data resource (http://www.perturbase.cn/). PerturBase is freely accessible, without any registration requirements.

## Supplementary data

Supplementary Data are available at NAR Online.

## Acknowledgements

We gratefully acknowledge all scPerturbation dataset owners for generously sharing their data. Special thanks to Eng. Qian Yu and Prof. Ling Guo for their invaluable technical support and insightful suggestions to enhance the database.

---

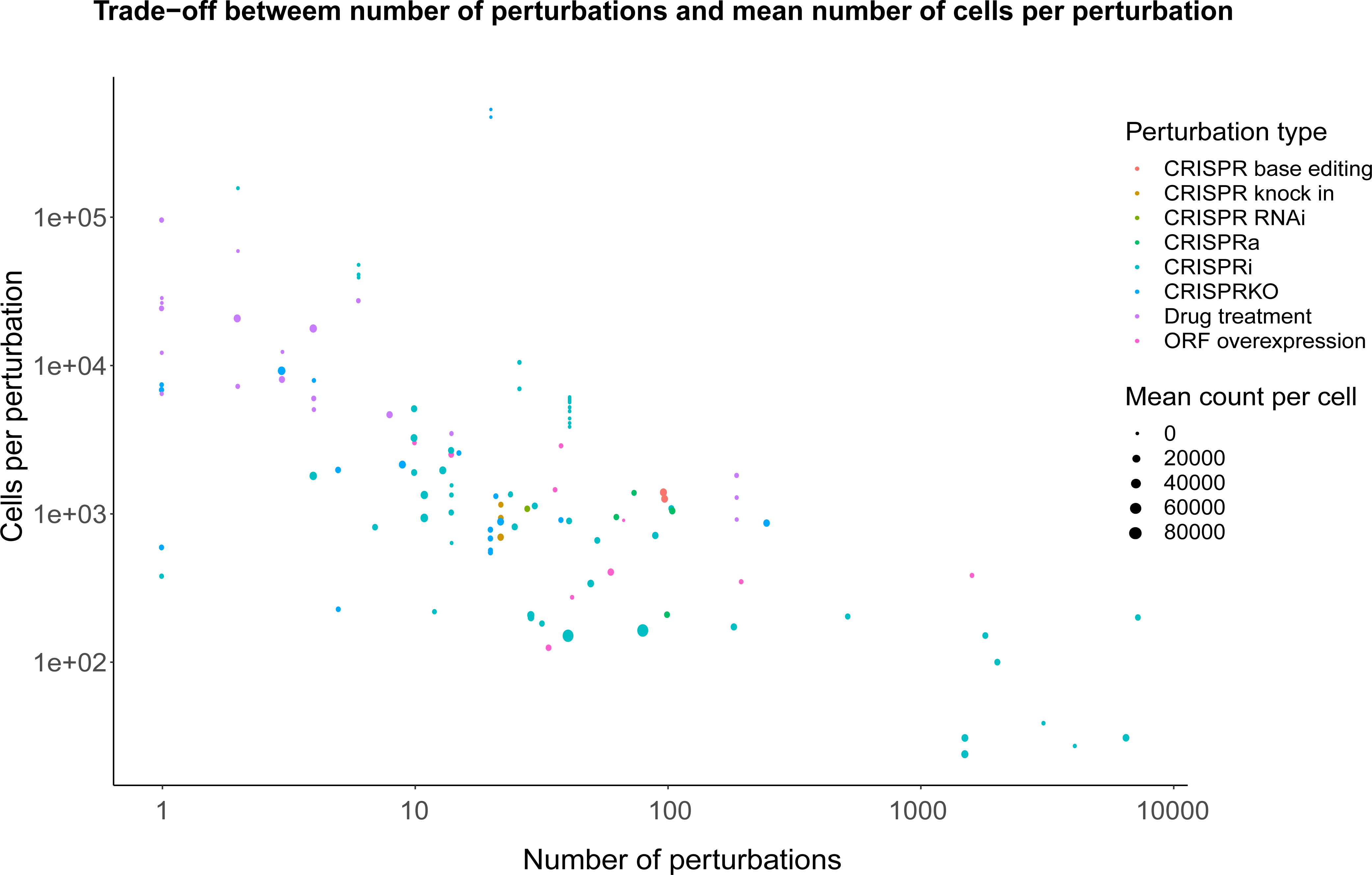

